# Ciliary photoreceptors in sea urchin larvae indicate pan-deuterostome cell type conservation

**DOI:** 10.1101/683318

**Authors:** Jonathan E. Valencia, Roberto Feuda, Dan O. Mellott, Robert D. Burke, Isabelle S. Peter

## Abstract

One of the signatures of evolutionarily related cell types is the expression of similar combinations of transcription factors in distantly related animals. Here we present evidence that sea urchin larvae possess bilateral clusters of ciliary photoreceptors that are positioned in the oral/anterior apical neurogenic domain and associated with pigment cells. The expression of synaptotagmin indicates that the photoreceptors are neurons. Immunostaining shows that the sea urchin photoreceptors express an RGR/G_O_-opsin, opsin3.2, which co-localizes with tubulin on immotile cilia on the cell surface. Furthermore, orthologs of several transcription factors expressed in vertebrate photoreceptors are expressed in sea urchin ciliary photoreceptors, including Otx, Six3, Tbx2/3, and Rx, a transcription factor typically associated with ciliary photoreceptors. Analysis of gene expression during sea urchin development indicates that the photoreceptors derive from the anterior apical neurogenic domain. Thus, based on location, developmental origin, and transcription factor expression, sea urchin ciliary photoreceptors are likely homologous to vertebrate rods and cones. However, we found that genes typically involved in eye development in many animals, including *pax6*, *six1/2*, *eya*, and *dac*, are not expressed in sea urchin ciliary photoreceptors. Instead, all four genes are co-expressed in the hydropore canal, indicating that these genes operate as a module in an unrelated developmental context. Thus, based on current evidence, we conclude that at least within deuterostomes, ciliary photoreceptors share a common evolutionary origin and express a shared regulatory state that includes Rx, Otx, and Six3, but not transcription factors that are commonly associated with the retinal determination circuit.

## INTRODUCTION

The remarkable similarity among cell types in distantly related animals suggests that some cell types have been present as a functional unit for a very long period of time during animal evolution [1–3]. The similarity between these cell types is reflected in specific structural and functional properties, and also at the molecular level in the expression of similar gene sets. In particular the expression of similar combinations of transcription factors is considered one of the signatures of cell type conservation [2].

A prominent example of a cell type that is broadly shared among metazoans are photoreceptor cells that are used for the detection of light [4]. Two types of photoreceptors are commonly encountered in bilateria. Ciliary photoreceptors are predominantly deployed in deuterostomes, while rhabdomeric photoreceptors are typically present in protostomes [5, 6]. However, ciliary photoreceptors are also present outside of deuterostomes, for example in the annelid *Platynereis dumerilii* [6] and in the cubozoan jellyfish *Triedalia cystophora* [7], while the presence of rhabdomeric photoreceptors has been shown in basal deuterostome sea urchins and in amphioxus [8–10]. Ciliary and rhabdomeric photoreceptors are morphologically distinct by being associated with different cell surface extensions that increase the photosensitive area. They are also molecularly distinct by deploying non-orthologous molecules for transducing the intracellular response to photo-excitement [11]. These molecular differences suggest that the two types of photoreceptors are non-homologous although they most likely co-existed in bilaterian ancestors [11]. It remains unclear however whether the two types of photoreceptors arose just once and subsequently gave rise to all the different variations of photoreceptors present in different animal clades, or whether photoreceptor cells evolved several times independently, by means of convergent evolution. To trace the evolutionary history of photoreceptor cell types within bilateria, data from different clades across the phylogeny are required.

Comparative analyses have demonstrated considerable similarity among the ciliary photoreceptors and eyes within vertebrates [12, 13]. At the morphological level, vertebrate eyes have a distinct position in respect to the brain, anterior and on the left and right side, and include rod and cone ciliary photoreceptors as well as neurons, pigment cells and other cell types. At the developmental level, eyes and photoreceptors derive from a similar embryological origin within vertebrates, the anterior neuroectoderm. Furthermore, ciliary photoreceptors in vertebrate eyes show similarities at the molecular level, expressing a similar type of c-opsins and a similar set of transcription factors including Otx, Rx, and Crx [14–20]. The similarity among eyes and photoreceptors extends to non-vertebrate chordates. Thus expression of Rx is required for the development of ocelli in *Ciona intestinalis* [21]. In amphioxus, the ciliary photoreceptors and pigment cells of the frontal eye show structural and molecular similarities with the ciliary photoreceptors and pigment cells of vertebrate eyes [8]. These similarities suggest that the ciliary photoreceptors in the frontal eye of amphioxus are homologous to the ciliary photoreceptors in vertebrate eyes, and accordingly, that ciliary photoreceptors descend from a common ancestral ciliary photoreceptor cell type, at least within chordates [2, 8]. However, the question remains whether the homology of ciliary photoreceptors extends to the entire deuterostome clade including basal deuterostome echinoderms.

Here we present evidence showing that the larvae of the purple sea urchin *Strongylocentrotus purpuratus* possess simple eyes consisting of ciliary photoreceptors and associated pigment cells. These photoreceptors are positioned on the left and right side of the bilateral larvae in the oral/anterior apical neurogenic organ with which they share a common developmental origin. In addition, the expression of several transcription factors including Rx, Otx, and Six3, but not Pax6, shows similarity to the regulatory state of vertebrate ciliary photoreceptors. These findings support the conclusion that the ciliary photoreceptors in echinoderms are homologous to the ciliary photoreceptors present in vertebrate eyes and thus descend from an ancestral ciliary photoreceptor cell type that was present in the last common ancestor of deuterostomes. Furthermore, the absence of pax6, six1/2, eya, and dac expression in the simple eyes of sea urchin larvae suggests that these genes might not have been part of the ancestral regulatory state of ciliary photoreceptors.

## RESULTS

### *Rx* and *opsin3.2* are co-expressed in bilateral clusters of cells

The transcription factor Rx plays an important role in vertebrate eye development [17]. Rx regulates expression of *pax6* and *six3* during early eye development, and the expression of *opsin* and other effector genes in differentiated ciliary photoreceptors in vertebrates [18, 20, 22, 23]. Rx is also expressed in ciliary photoreceptors of non-vertebrate chordates, including amphioxus [8] and *Ciona intestinalis* [21], and in protostomes, such as the marine annelid *Platynereis dumerilii* [6]. These observations suggest that the requirement of Rx in the formation of ciliary photoreceptors is broadly shared among bilaterian animals. We thus examined the expression of *rx* in larvae of a non-chordate basal deuterostome, the purple sea urchin *Strongylocentrotus purpuratus*. When analyzed by whole mount in situ hybridization (WMISH), expression of *rx* was restricted to bilateral clusters of 2-3 cells located on the oral side of the larva between mouth and the neurogenic apical organ (Fig. 1A). The particular location of these cells, and the expression of *rx*, suggested that these cells might correspond to photoreceptor cells. To test this hypothesis, we analyzed other features that are typically associated with photoreceptors: i) the expression of a photosensitive opsin, ii) the potential to elicit a neuronal response, and iii) the presence of shading pigments.

**Figure 1.**
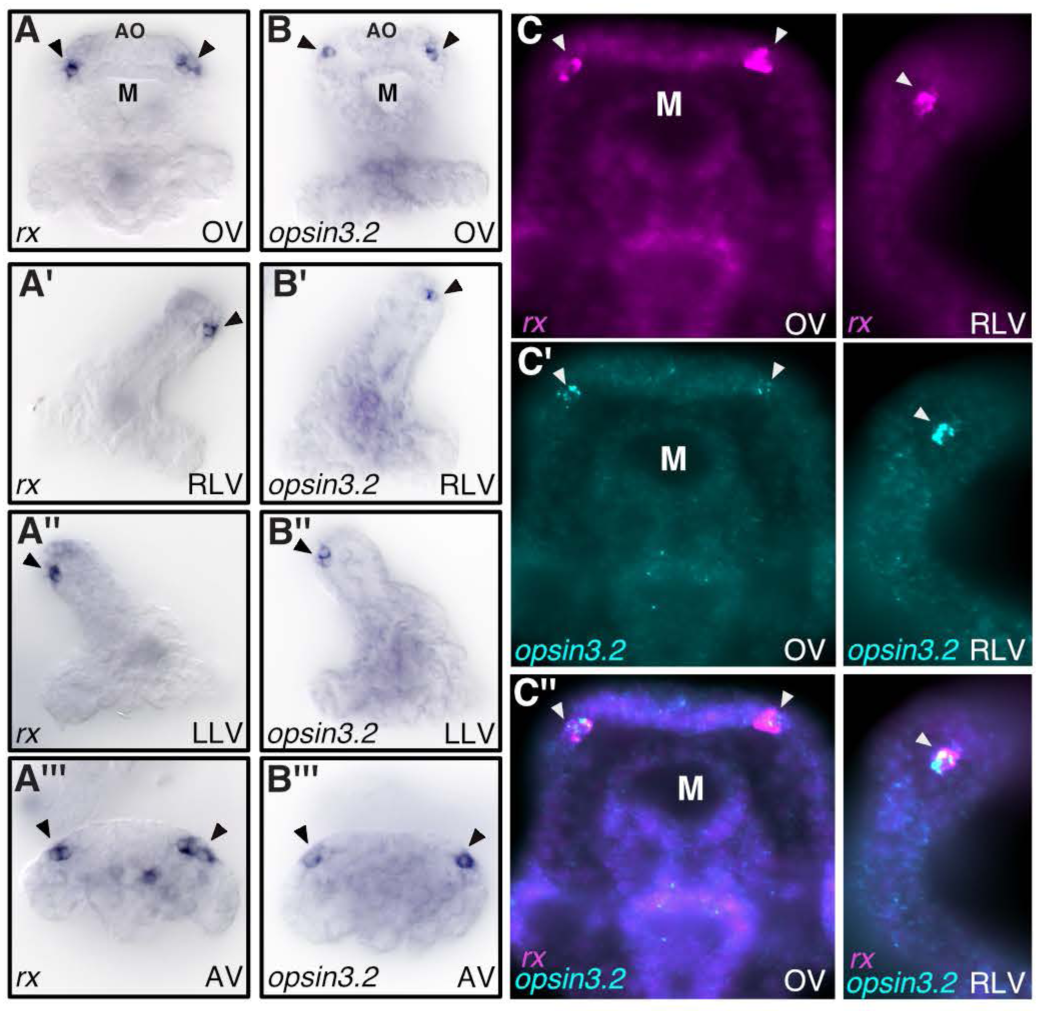
Spatial expression of *rx* and *opsin 3.2* in putative photoreceptors of sea urchin larvae. **(A-A’’’)** WMISH showing *rx* expression in 72h larvae on oral side of apical organ. **(B-B’’’)** WMISH for *opsin3.2* expression. **(C-C’’)** Double fluorescent WMISH for *rx* (magenta, **C**) and *opsin3.2* (green, **C’**), overlay shown in **C’’**. Arrowhead indicates photoreceptors. M, mouth; AO, apical organ; OV, oral view; RLV, right lateral view; LLV, left lateral view; AV, apical view.

To determine if the putative larval photoreceptor cells express opsins, available transcriptome data were analyzed for the presence of *opsin* transcripts during sea urchin development [24]. Eight opsin genes are encoded in the sea urchin genome, of which only *opsin2*, and *opsin3.2* showed expression at the larval stage (72h), while transcripts of other opsin genes were not detectable during early development (Fig. S1A). To determine whether these *opsin* genes are expressed in putative photoreceptors, we analyzed the spatial expression of *opsin2* and *opsin3.2* by WMISH. The results show that *opsin2* is expressed in the stomodeum (mouth) region and part of the ciliary band between the lower arms (Fig. S1B). On the other hand, *opsin 3.2* is expressed in bilateral clusters of cells similar to *rx*, consistent with a previous study (Fig. 1B and Fig. S1B) [25]. We performed double fluorescence WMISH to determine whether *rx* and *opsin3.2* are co-expressed in the same cells. The results confirm that *rx* and *opsin3.2* are both co-expressed in the putative photoreceptors (Fig. 1C). A phylogenetic analysis showed that opsin3.2 belongs to the RGR/Go class (or Group 4) of opsins (Fig. S2), the same class of opsins that are also expressed in ciliary photoreceptors of scallops [26–28] and in rhabdomeric photoreceptors in *Platynereis* [29]. According to this analysis, opsin3.2 is a co-ortholog of *Platynereis* opsin1, which is sensitive to cyan light at wavelengths important for marine life [29].

### Putative photoreceptors express synaptotagmin

The response to light is transmitted from photoreceptors to the nervous system through synaptic exocytosis of neurotransmitters [30]. The bilateral clusters of *opsin3.2* expressing cells in the sea urchin larva are positioned adjacent to the neurogenic apical organ. To determine if the putative photoreceptor cells possess synapses, we analyzed the expression of synaptotagmin, a pre-synaptic neuronal protein involved in vesicle exocytosis [30–32]. Immunostaining was performed with antibodies against the pan-neuronal synaptotagmin B [33] and with rat polyclonal antibodies against sea urchin opsin3.2 in 72h sea urchin larvae. Expression of opsin3.2 protein was detected in bilateral clusters of cells, similar to the localization of *opsin3.2* mRNA (Figs 2A and 2B). The results show that opsin3.2 is expressed in a subset of synaptotagmin B expressing cells (Figs 2A, 2C, Fig. S3A). This result shows that the putative photoreceptor cells are neuronal. In comparison, co-immunostaining of opsin3.2 and serotonin showed expression in separate cells, indicating that, as expected, the photoreceptor cells are not serotonergic neurons (Fig. S3B).

**Figure 2.**
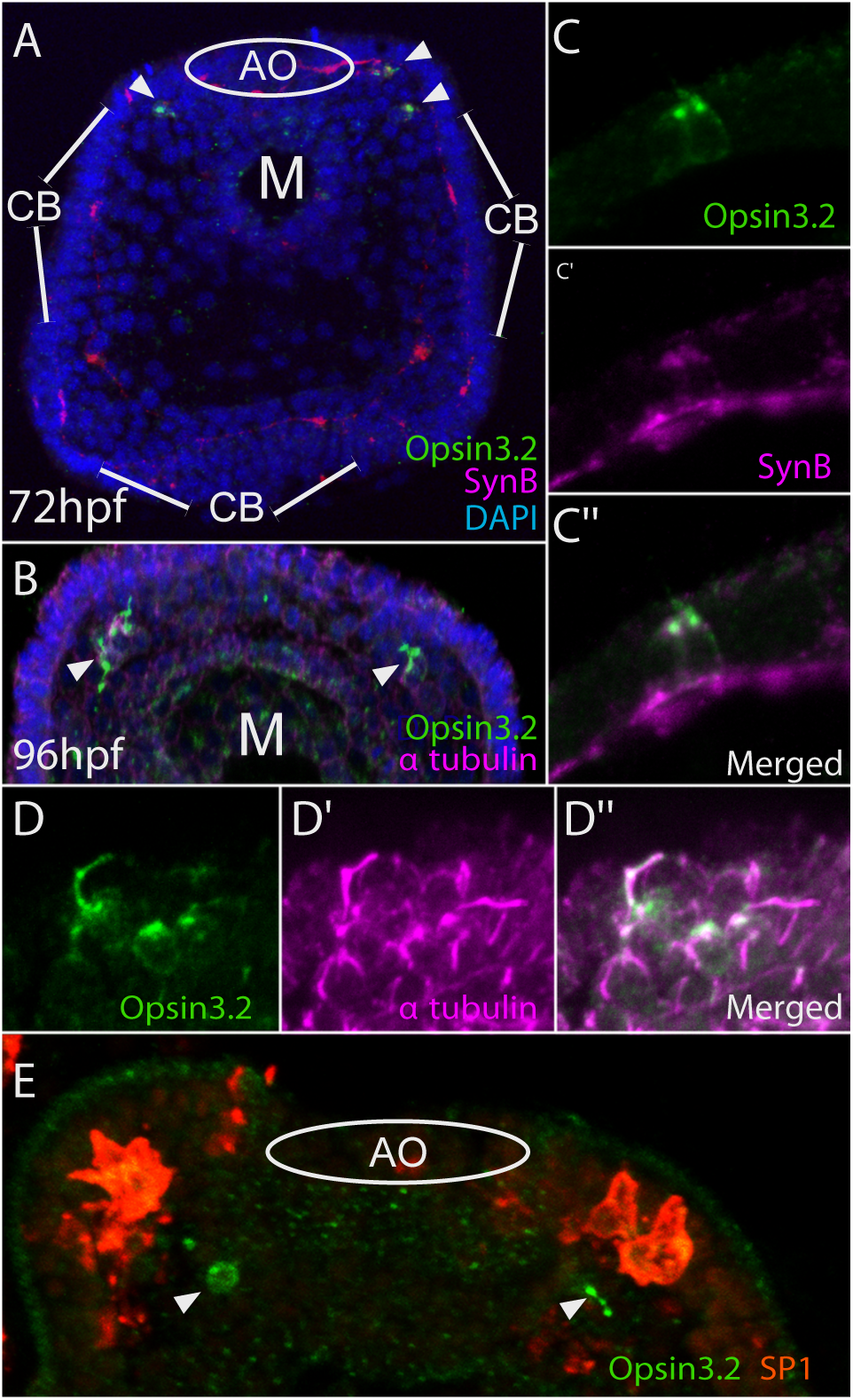
Immunostaining showing opsin 3.2 expression in neuronal ciliary photoreceptors. Confocal laser scanning images of whole mount *S. purpuratus* larvae showing immunostaining **(A)** of opsin 3.2 and synaptotagmin B at 72h, and **(B)** of opsin3.2 and *α* tubulin at 96 h. **(C)** Confocal images of co-immunostaining for opsin3.2 (green, **C**) and synaptotagmin B (magenta, **C’**), showing co-expression (**C’’**). **(D)** Confocal images of co-immunostaining for opsin3.2 with *α*-tubulin **(D’)** showing co-localization in cell surface cilia of photoreceptors **(D’’)**. **(E)** Co-immunostaining of opsin3.2 (green) and pigment cell specific SP1 (red) showing pigment cells located in proximity to photoreceptors. M, mouth; AO, apical organ; CB, ciliated band.

### *Rx* expressing cells correspond to ciliary photoreceptors

Ciliary and rhabdomeric photoreceptor cells possess morphologically distinct cell surface structures that increase the photosensitive area [34]. The expression of *rx* suggested that the sea urchin larval photoreceptors belong to the class of ciliary photoreceptors. We thus determined whether the photoreceptor cells are equipped with immotile cell surface cilia. We used antibodies against *α*-tubulin to detect the presence of microtubules, a structural component of cilia, on the surface of the sea urchin photoreceptor cells. The results show that opsin3.2 and *α*-tubulin co-localize on short cilia that are present on the apices of photoreceptor cells (Figures 2B and 2D). The short cilia could be identified in living specimens and appear to be immotile. Taken together, the presence of opsin presenting immotile cilia on the surface of Rx expressing neurons support the conclusion that the bilateral clusters of cells in sea urchin larvae correspond to ciliary photoreceptors.

### Ciliary photoreceptors are associated with pigment cells

In order to enable a directional perception of light, photoreceptors are typically associated with shading pigments [11, 34]. To determine if shading pigments are present near the photoreceptors in sea urchin larvae, immunostaining was performed in 72h larvae using SP1 antibodies to detect pigmented immunocytes [35]. Indeed, pigment cells were found within 2-3 cell diameters of the photoreceptors, embedded in the ectodermal epithelium (Fig. 2E). Although the pigment cells are not in immediate contact with the photoreceptor cells, they still potentially provide sufficient shading to enable directional photoreception. In addition, pigment cells are dispersed throughout the aboral ectoderm, but completely absent from the oral ectoderm, potentially further biasing the intensity of light perceived from the oral versus aboral side of the larva.

### Transcription factor expression and developmental origin of ciliary photoreceptors

Several transcription factors were found in addition to Rx to be expressed in sea urchin ciliary photoreceptors at 72h: *arrowhead* (*Sp-awh*; Lhx6/8-like), *forkhead box G* (*foxg*), *homeobrain* (*hbn)*, *nk2 homeobox 1 (nkx2.1), otx*, *sine oculis-related homeobox 3* (*six3)*, *soxb2*, *T-box 2/3* (*tbx2/3*), and *zic* (Fig. 3A and Fig. S4). In addition, expression of the *inhibitor of DNA binding* transcription factor *Id* was detected at 60h and weakly also at 72h in photoreceptors (Fig. 3 and Fig. S4). To test whether these regulatory genes are expressed in photoreceptor cells, expression of *awh* and *six3* was furthermore analyzed by double fluorescent WMISH together with probes detecting *opsin3.2*. The results showed for both genes overlapping expression with *opsin3.2*, confirming expression in photoreceptor cells (Fig. S5). Several of the transcription factors expressed in sea urchin photoreceptors are also expressed in vertebrate eyes and photoreceptors, as we will discuss below.

**Figure 3.**
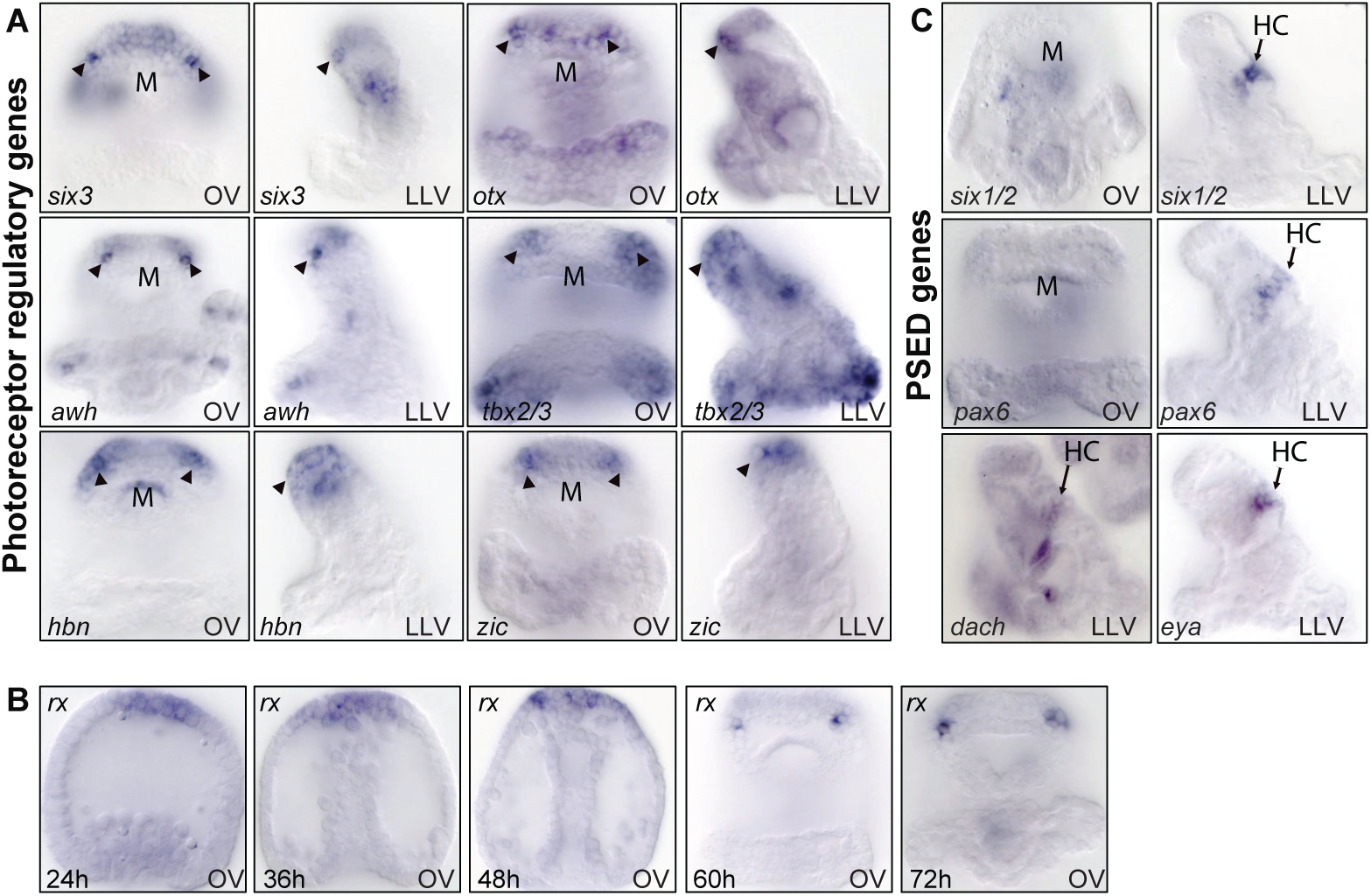
Expression of photoreceptor regulatory state and PSED genes in sea urchin larvae. **(A)** WMISH in 72h larvae showing expression of regulatory genes in ciliary photoreceptors (arrowheads). Additional regulatory genes expressed in photoreceptors are shown in Figure S4. **(B)** WMISH in 72h larvae showing expression of *pax6*, *six1/2*, *dach*, and *eya* in the hydropore canal (HC). **(C)** Developmental time course of *rx* expression indicating specification of progenitors of photoreceptors in apical neurogenic domain. OV, oral view; LLV, left lateral view.

To determine the developmental time course for photoreceptor specification, gene expression was analyzed at several stages between 24-72h. Based on transcriptome time course data, expression of *opsin3.2* is initiated at 60h, indicating that the differentiation of photoreceptor cells occurs sometime after 60h (Fig. S1A) [24]. Furthermore, we analyzed the expression of *rx* and other regulatory genes at several stages of development by WMISH (Fig. 3B and Fig. S4). The results show that *rx* is expressed broadly in the apical neurogenic domain at 24h, as shown earlier [36], and expression becomes restricted in the oral/anterior region of the apical domain to the photoreceptors by 60h (Fig. 3B). This expression pattern shows similarity to expression of *rx* in vertebrates. In mice for instance, *rx* expression initiates in the anterior neural plate before becoming progressively restricted to eyes and ventral forebrain, and later to photoreceptors [17]. Similarly, all transcription factors examined here are expressed in the oral apical domain by 36h. Of these, *awh* and *six3* show specific expression in photoreceptors within the oral apical domain by 60h, while the other transcription factors are expressed in broader areas including the photoreceptors (Fig. 3B and Fig. S4). These results show that the expression of photoreceptor-associated transcription factors occurs in the anterior/oral region of the neurogenic apical domain before the differentiation of photoreceptors and expression of opsins. Based on position and expression of transcription factors, these results suggest that the anterior neurogenic region provides the developmental origin of the larval photoreceptors. Photoreceptor precursors originating in the oral apical neurogenic domain become distinctly specified as photoreceptors by 60h, when expression of *opsin3.2* is initiated.

### Expression of *pax6*, *six1/2*, *dach*, and *eya* in the hydropore canal but not in ciliary photoreceptors

Given their important role in eye development and the specification of retinal progenitor cells throughout bilateria, we analyzed the expression of *pax6*, *six1/2*, *eya*, and *dach* (PSED genes) in 72h sea urchin larvae by WMISH (Fig. 3C). Surprisingly, the expression of all four regulatory genes was detected in the hydropore canal, a mesodermal derivative for filtering and secretion of the coelomic fluid. However, the expression of these PSED genes was absent from larval photoreceptors at 72h (Fig. 3C and Fig. S6) [37, 38]. To analyze whether expression of these genes occurred during earlier stages of photoreceptor specification, *pax6*, *six1/2*, and *eya* expression was analyzed at 24h, 36h, 48h, and 60h (Fig. S6). Our results indicate that the PSED module does not operate during development or differentiation of the sea urchin ciliary photoreceptors, even though a functional PSED module is encoded in the genome and expressed in the hydropore canal.

## Discussion

### Presence of photoreceptors in echinoderms

The results shown here provide evidence that sea urchin larvae possess bilateral clusters of ciliary photoreceptors that are positioned at the oral side of the larvae, between nervous system and mouth. These photoreceptors express opsins that are present in particular on immotile surface cilia. The association of these ciliary photoreceptors with pigment cells suggests that sea urchin larvae possess simple eyes that are capable of directional light perception [4]. Interestingly, the tube feet of adult animals of the same sea urchin species were previously shown to include rhabdomeric photoreceptors, based on their expression of opsin 4 and Pax6 [10]. Thus, just like amphioxus, *Platynereis* and several other species, sea urchins possess both ciliary and rhabodomeric photoreceptors. In sea urchins, these photoreceptors are present at different stages in the life cycle. Only ciliary photoreceptors are present in bilateral pluteus larvae, at least at the stages considered here. However, the expression of molecular markers including opsins and Pax6 in the tube feet of adult sea urchins suggests that both ciliary and rhabdomeric photoreceptors are present in penta-radial adult animals [10, 39].

### Expression of an atypical opsin in sea urchin ciliary photoreceptors

The ciliary photoreceptors in most deuterostome species and also a few protostomes typically express c-opsins. However, despite the expression of a set of transcriptional regulators similar to vertebrate photoreceptors, sea urchin ciliary photoreceptors express a non-typical type of opsin, an RGR/Go opsin. Members of the RGR/Go family of opsins are expressed in ciliary and rhabdomeric photoreceptors in several protostome species. Although the RGR/Go class of opsins is used less frequently, a member of this class is for example expressed in the ciliary photoreceptors of scallops [4, 28]. Based on the phylogenetic distribution of c-opsin expression in ciliary photoreceptors in bilateria, it seems likely that expression of c-opsin represents the ancestral state in ciliary photoreceptors while the co-option of RGR/Go opsins might have occurred within the echinoderm lineage. Similar co-options of different types of opsin presumably occurred relatively frequently during photoreceptor evolution [13].

### Evidence for homology of deuterostome ciliary photoreceptors

Two types of evidence suggest that the ciliary photoreceptors described here are homologous to the ciliary photoreceptors present in vertebrates and other deuterostome species: the expression of a common set of transcription factors and a similar developmental origin. Thus, the combination of transcription factors expressed in the sea urchin ciliary photoreceptors show a molecular signature that is similar to other deuterostome ciliary photoreceptors (Figures 4A and 4B). Most prominently, Rx has been shown to be expressed in ciliary photoreceptors of many species, including amphioxus, *Ciona*, *Platynereis*, and vertebrates [8, 14, 17, 21, 23, 40]. In addition, the sea urchin photoreceptors express Otx, an ortholog of Otx2 and Crx that are involved in photoreceptor specification and differentiation in vertebrates [13]. Otx is also involved in photoreceptor specification of amphioxus [8] and brachiopods [41]. Six3 is broadly expressed in anterior neural plate regions including photoreceptors of amphioxus [8], while its ortholog Six7 is required for the development of photoreceptors and expression of opsin in zebrafish [42, 43]. Six3 is also expressed in the mouse retina and has been associated with the regulation of rhodopsin expression [44]. Furthermore, members of the Tbx2 family are involved in the specification of UV-cone cells in the zebrafish retina [45] and in the specification of the eye field in Xenopus [22]. Expression of Id is detected at 60h in sea urchin photoreceptors, but only weakly at 72h. In vertebrates, orthologs of Id transcription factors are expressed in retinal progenitor cells in zebrafish and mice and are downregulated in differentiating photoreceptors [46, 47]. Zic orthologs are expressed in retinal progenitor cells in mice [48].

**Figure 4.**
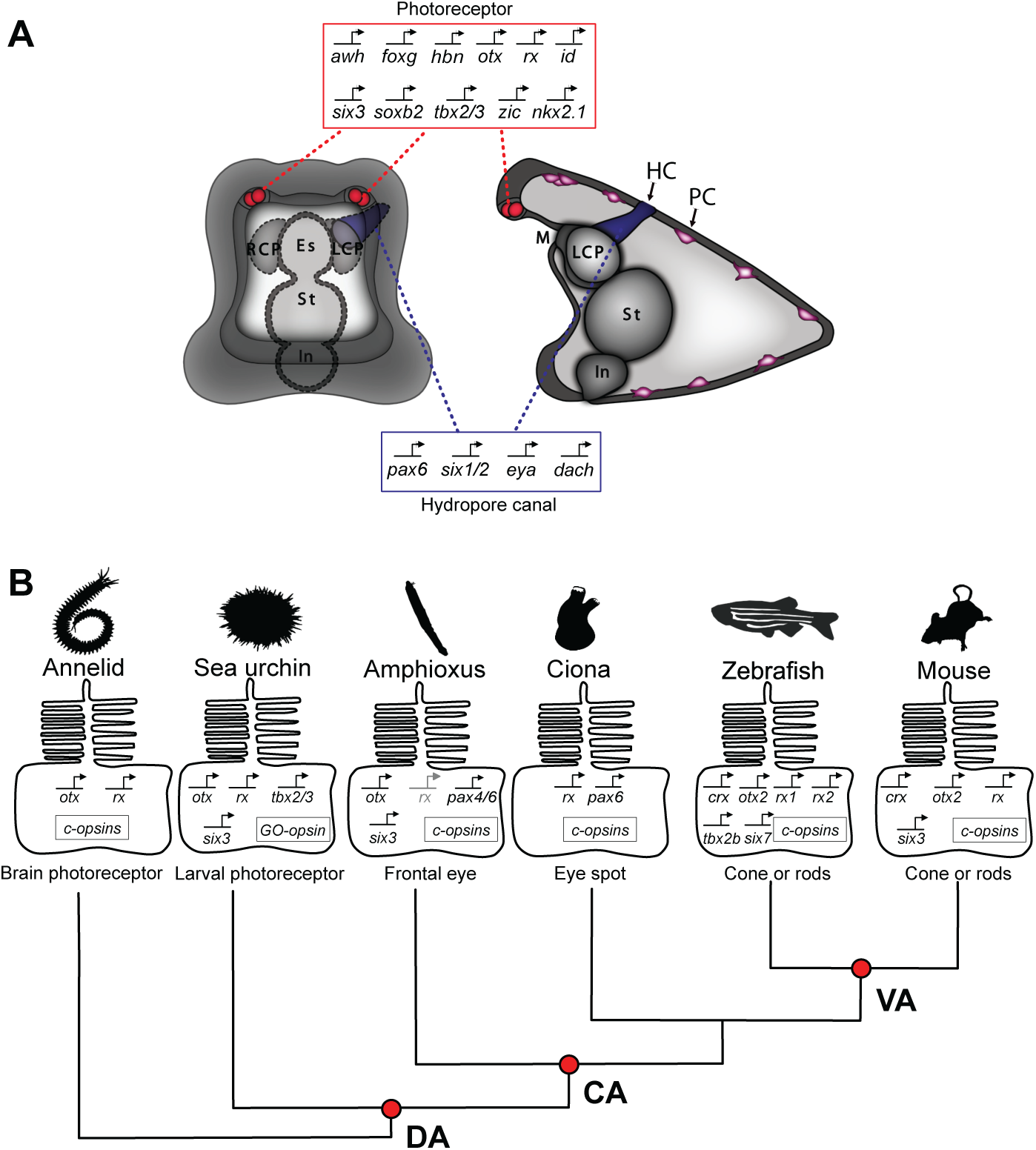
Summary diagram showing sea urchin photoreceptors and phylogenetic distribution of photoreceptor-specific regulatory states. **(A)** Summary diagram of sea urchin larva showing expression of regulatory genes in photoreceptors and expression of indicated PSED genes in the hydropore canal (HC). Pigment cells (PC) are associated with photoreceptors and distributed throughout aboral ectoderm. **(B)** Diagram showing expression of a common set of regulatory genes in ciliary photoreceptors in animals representing specific positions within the phylogenetic tree. DA, deuterostome ancestor; CA, chordate ancestor; VA, vertebrate ancestor.

Thus, as summarized in Fig. 4B, these data indicate that there is a common regulatory toolkit used for the differentiation of ciliary photoreceptors, including Rx, Otx and its ortholog Crx, Six3, and possibly Tbx2. Thus, a significant fraction of the regulatory state expressed in sea urchin larval photoreceptors is expressed also in ciliary photoreceptors in other deuterostomes as well as in protostome ciliary photoreceptors. Including Zic and Id that are expressed in retinal progenitors, over half of the sea urchin photoreceptor transcription factors described here are involved in eye development in vertebrates. These results therefore suggest that ciliary photoreceptors express a conserved cell-type specific regulatory state that already contributed to the specification of ancestral ciliary photoreceptors.

Furthermore, gene expression data suggest that sea urchin larval photoreceptor cells developmentally derive from cells of the oral/anterior region of the apical neurogenic ectoderm. The developmental progression of *rx* expression, for example, is at first broadly distributed in the anterior neural ectoderm, and later progressively restricted to the photoreceptor cells in sea urchin larvae and similarly also in many vertebrate species [14]. The developmental formation of sea urchin photoreceptors is also reminiscent of the neural ectodermal origin of photoreceptors in *Ciona* and amphioxus [49]. Together, the similarity in transcription factor expression and embryonic origin of sea urchin ciliary photoreceptors suggest that ciliary photoreceptors are evolutionarily homologous at least within deuterostomes.

### Pax6 and the evolutionary history of eyes and ciliary photoreceptors

A remarkable observation that puzzles evolutionary biologists is that transcription factors related to Pax6, Six1/2, Eya, and Dach (PSED), that compose the retinal determination network, are involved in eye development in Drosophila, vertebrates and many other animals [50]. The important role of Pax6 in particular has been demonstrated in many animals where mutation of *pax6* typically leads to severe effects in eye development. The extraordinary similarity in the requirement of Pax6 during early eye development in distantly related animals including Drosophila [51] and vertebrates [52], and the ability of Pax6 to induce ectopic eyes when overexpressed in ectopic locations [53, 54], has led to the hypothesis that Pax6 and several other transcription factors of the retinal determination network function as “eye master-regulators” that were expressed in the proto-eyes of bilaterian ancestors [55]. However, such apparent molecular homology is in sharp contrast to the conclusions of comparative analyses of eye development and morphology, which indicate that complex eyes evolved independently several times during bilaterian evolution. The question therefore arises whether the expression of a shared set of transcription factors in the eyes of flies and vertebrates is a result of evolutionary conservation and already present in bilaterian ancestors, or a result evolutionary co-options of PSED regulatory circuits during the evolution of complex eyes [56].

In order to address this question, it becomes important to determine whether the absence of PSED factors in sea urchin ciliary photoreceptors reflects an evolutionary loss of function that occurred specifically in the echinoderm lineage, or whether the sea urchin photoreceptors likely represent the ancestral state of simple eyes in deuterostome ancestors. To reconstruct the most likely ancestral state of ciliary photoreceptors within deuterostomes, a comparison with other deuterostome species becomes necessary. Pax6 for example although not expressed in sea urchins, is expressed in the ciliary photoreceptors of invertebrate chordates such as amphioxus and *Ciona* [8, 57]. In vertebrates, Pax6 is specifically required during early eye development and contributes to the maintenance of multi-potency in retinal progenitor cells [52, 58]. However, during later stages of retinal development, Pax6 controls the specification of horizontal and amacrine cells and inhibits the differentiation of ciliary photoreceptor cells [59]. Thus Pax6 functions in the progenitors of photoreceptors but not in differentiated ciliary photoreceptors. This is consistent with the observation that expression of differentiation genes such as *c-opsins* does not require Pax6 in vertebrate ciliary photoreceptors, but is controlled by Otx and Rx [14, 60]. The absence of Pax6 function in ciliary photoreceptors of echinoderms and vertebrates is not consistent with the view that Pax6 constitutes a crucial and conserved component of the cell type-specific regulatory state in ciliary photoreceptors. It appears more likely that Pax6 was not expressed in the ciliary photoreceptors of deuterostome ancestors but has been co-opted independently in non-vertebrate chordates. The scenario seems to be even more evident for other PSED transcription factors. Thus in amphioxus, PSED factors are co-expressed in many cell types but not in ciliary photoreceptors [61]. And while several PSED transcription factors are expressed during early eye development in vertebrates, their function, in particular during the differentiation of photoreceptors, is not as clearly resolved as in Drosophila [56, 60]. The absence of PSED transcription factor expression in sea urchin and amphioxus ciliary photoreceptors suggests that the PSED module was not required for the specification of ciliary photoreceptors in deuterostome ancestors and must have been co-opted during the evolution of the complex eyes of vertebrates.

Consistent with the idea that PSED transcription factors might have been co-opted to eye development during chordate evolution, PSED genes have been co-opted as a module to many other developmental processes. Thus, in vertebrates, PSED factors contribute to kidney development and the specification of somitic muscle [62], and in amphioxus, PSED factors are co-expressed in several cell fates other than photoreceptors [61]. Similarly, PSED genes are expressed in coelomic pouches of *Lytechinus variegatus* sea urchins [38] and, as we show here, the larval hydropore canal in *S. purpuratus*. The PSED module therefore must control functions that are required in many different developmental contexts, and has possibly been co-opted to control early eye development in vertebrates [63].

The evidence presented here suggests that, based on developmental and molecular similarities, ciliary photoreceptors in basal deuterostome sea urchins are homologous to ciliary photoreceptors that are present in chordates. Thus deuterostome eyes consist of a conserved ciliary photoreceptor cell type expressing a conserved regulatory state and sharing a common evolutionary origin in the eyes of deuterostome ancestors. Many subsequent evolutionary modifications must have occurred in the gene regulatory network controlling eye development, possibly including the co-option of the PSED circuit, to give rise to the variety of photoreceptors and eyes within deuterostomes.

## Materials and Methods

### Phylogenetic analysis and identification of orthologs

Opsin dataset was obtained by merging the sequences from [29] and [27]. Furthermore, additional opsin genes were obtained from the genomes of *Branchiostoma floridae* [64], *Branchiostome belechei* [65], *Ciona intestinalis* [66] and *Ciona savignyi* [67]. Specifically, the dataset of [26] composed by 449 sequences was used as seed and potential homologs were identified using BLASTP [68]. Each sequence with a e-value < 10^−10^ was retained a good opsin homolog. To identify opsin genes, sequences were further annotated using interproscan [69], and only sequences with retinal binding domains were considered as Opsins. The final dataset includes 232 Opsins and 10 melatonin genes that have been used to root the trees. Alignment was performed using MAFFT [70] and phylogenetic reconstruction was performed under Maximum likelihood framework and Bayesian framework under LG-G_4_ [71]. The ML tree was reconstructed using iqtree [72] and nodal support was estimated using ultrafast bootstrap [73] (1000 replicates) and the SH-aLTR bootstrap [74]. Bayesian inference was performed using Phylobayes4.1 [75] with two independent runs. Convergence was evaluated using tracecomp and bpcomp packages in Phylobayes (see Phylobayes manual). Alignment and trees are available at https://github.com/RobertoFeu/Opsins_phylogeny_Valencia_et_al. The identification of orthologs in Fig. 4B was performed using EggNOG mapper [76].

### Gene amplification and probe synthesis

The primer sets used for gene amplification are listed in TableS1. Gene models generated from sea urchin transcriptome analysis were used as a reference for primer design [24] using T7 tailed primers or cloning. cDNA prepared from various developmental stages was used as template for PCR. For cloning, PCR products were purified and ligated into GEM-T EZ constructs. Cloned genes were PCR-amplified using the primer flanking the insert region, and PCR products were used to synthesize RNA probes for WMISH.

### Whole-Mount in Situ Hybridization

The protocol for whole-mount *in situ* hybridization (WMISH) to detect spatial gene expression has been described previously [77]. Briefly, sea urchin embryos were fixed in 4% paraformaldehyde solution. The fixed embryos were incubated in hybridization buffer [50% (vol/vol) formamide, 5× SSC, 1× Denhardt’s, 1 mg/mL yeast tRNA, 50 ng/mL heparin, and 0.1% tween-20] with a concentration from 1 to 2 ng/µL digoxygenin RNA probe(s) at 60 °C for 18 h. Two Post hybridization washes were performed with hybridization buffer without RNA probe, 2× SSCT (2× SSC, 0.1% tween-20), 0.2× SSCT, and 0.1× SSCT, each 20 min at 60 °C. Subsequently, 5 washes were performed with a buffer of 0.1% Tween 20, 10% MOPS (1M), 10% NaCl (5M) and 80% DEPC water. Antibody incubations were performed at room temperature with 1:2,000 diluted anti-DIG Fab (Roche). The embryos were extensively washed before staining reaction, including six times with MABT buffer (0.1 M maleic acid, 0.15 M NaCl, and 0.1% tween-20), twice with AP buffer [100 mM Tris·Cl (pH 9.5), 100 mM NaCl, 50 mM MgCl2, and 1 mM levamisole]. 5-Bromo-4-chloro-3-indolyl-phosphate (BCIP) and nitro blue tetrazolium were used for staining. Fluorescent in situ *in situ* hybridization protocol was performed as described in [78].

### Antibody production

Antibody production was as previously described [79]. Antigens were made using a pET28b (+) plasmid (Novagen) for expression of 6XHis tagged proteins. An Opsin3.2 construct was prepared with PCR (opsin-cyto:F = 5’-CAGTCATATGGCGTCGGTAAAATAAG-3’, opsin-cyto:R = 5’-AGTCAAGCTTCTGTAGATTTTTAATG-3’) encoding the carboxyl cytoplasmic domain (844-1494 of the coding sequence and 282-498 of the protein). High fidelity PCR was used with a cDNA template prepared from *S. purpuratus* embryos and the 650 bp product was cloned using the pGEM-T Easy system (Promega). Protein expression was induced in E. coli (BL21). Bacterial lysate was prepared and protein was solubilized in binding buffer (6 M guanidine HCl, 0.5 M NaCl, 100 mM Na2HPO4, 100 mM NaH2PO4, 10 mM imidazole, 10 mM Tris, 1 mM 2-mercaptoethanol, pH 8.0) prior to affinity purification by immobilized metal ion affinity chromatography (IMAC) using Chelex 100 Resin (Bio-Rad). Purified protein in PBS was mixed 1:1 with Freund’s complete adjuvant for immunization or with Freund’s incomplete adjuvant for booster injections. A rat was immunized by subcutaneous injection of 100 mg antigen in 250 µl of adjuvant, and booster injections were done 21 days and 42 days after the initial immunization. Terminal bleed via cardiac puncture was done after 52-56 days. Blood was incubated at 37°C for 45 min and then 4°C overnight. Samples were centrifuged at 1,000 XG, and serum collected. Antibody specificity was established by pre-absorbing the immune serum with an approximately equimolar preparation of the protein used to immunize the rat. Pre-absorption eliminated antibody binding to 72 and 96 h larvae.

### Immunofluorescence

*S. purpuratus* embryos were collected at the desired time point and fixed for 5-10 min in 4% paraformaldehyde in PEM buffer [80]. Embryos were washed with phosphate buffered saline (PBS), blocked for 1 h in SuperBlock (Thermo), probed with primary antibody, and washed 3 times with PBS. Alexa Fluor fluorescent secondary antibodies (Invitrogen) were used to visualize antibody labeling on a Zeiss 700 LSM (Carl Zeiss) confocal microscope. All preparations were done at 4° C. Imaging and analysis was conducted using ZEN (2009) or ImageJ (1.44) software. Adobe Photoshop (9.0.2) was used to prepare figures and adjust image contrast and brightness. Antibodies employed anti-SynB [33]; Sp1 [35]; a-tubulin (Santa Cruz Biotechnologies, sc-23948).

## Acknowledgements

We thank Drs. Markus Meister and Daniel Wagenaar for helpful discussions on earlier versions of the manuscript. This work was supported by National Institutes of Health Grants HD 037105 and HD 094047 (to I.S.P.), and by a Discovery Grant from the Natural Sciences and Engineering Research Council of Canada (2016-03737), awarded to R.D.B.

## Author contributions

The project was initially conceived by J.V., R.F., and I.S.P; J.V. and R.F. performed WMISH analysis; D.O.M. and R.D.B. contributed antibodies and performed immunostaining; J.V., R.F. and I.S.P. contributed analysis of PSED circuit in the context of GRN evolution; R.F., R.D.B., and I.S.P. wrote the manuscript.

## SUPPLEMENTAL INFORMATION

**Figure S1.**
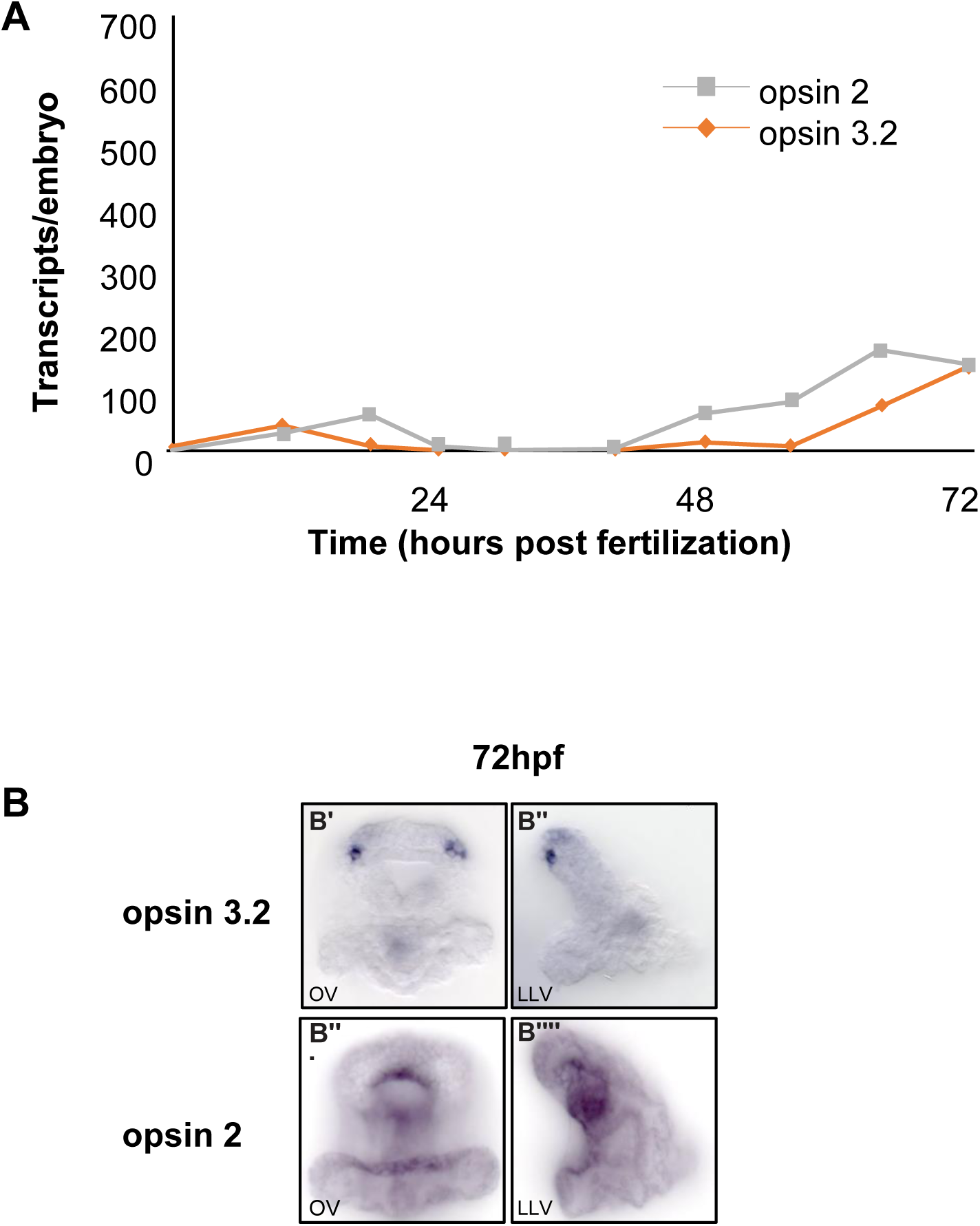
Expression of opsin genes in putative photoreceptors in sea urchin larvae. **(A)** Developmental time course of opsin gene expression based on transcriptome data from [1]. **(B)** Spatial expression of opsin 3.2 and opsin 2 in 72h sea urchin larvae. OV, oral view; LLV, left lateral view.

**Figure S2.**
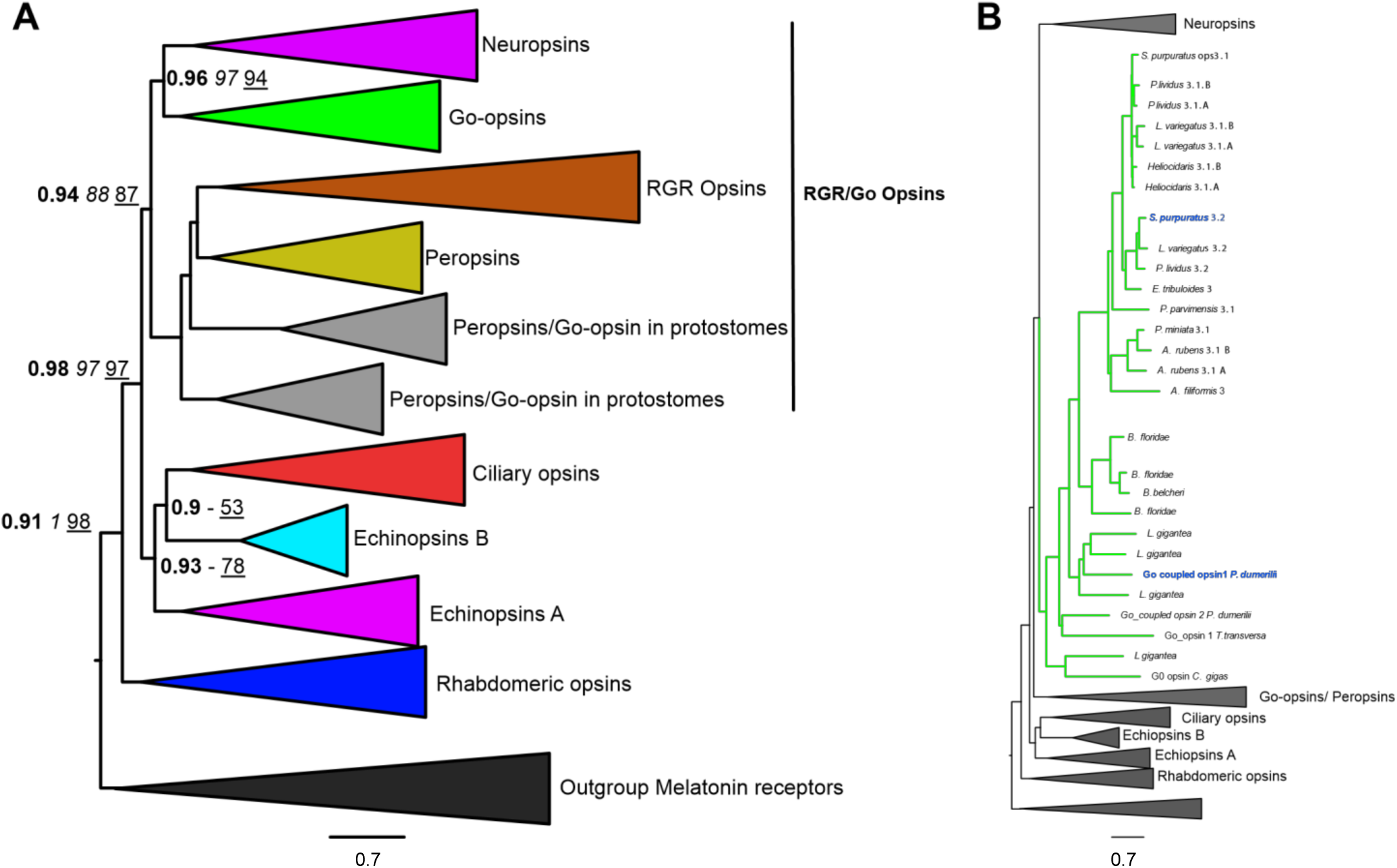
Phylogenetic analysis of opsins. (A) Phylogenetic tree of opsins. Values in bold indicate the Bayesian posterior probability, values in italics indicate SH-aLTR bootstrap values, values underlined indicate ultrafast bootstrap (1000 replicates). Majore opsin clades are color coded. The three group-topology where R-opsins are a sister group of C-opsins and RGR/Go-opsins is well supported [2]. (B) Phylogenetic tree as in (A) with focus on Go-opsins. Opsin 3.2 is a co-ortholog of Go-opsin1 of *Platynereis dumerilii* [3].

**Fig. S3.**
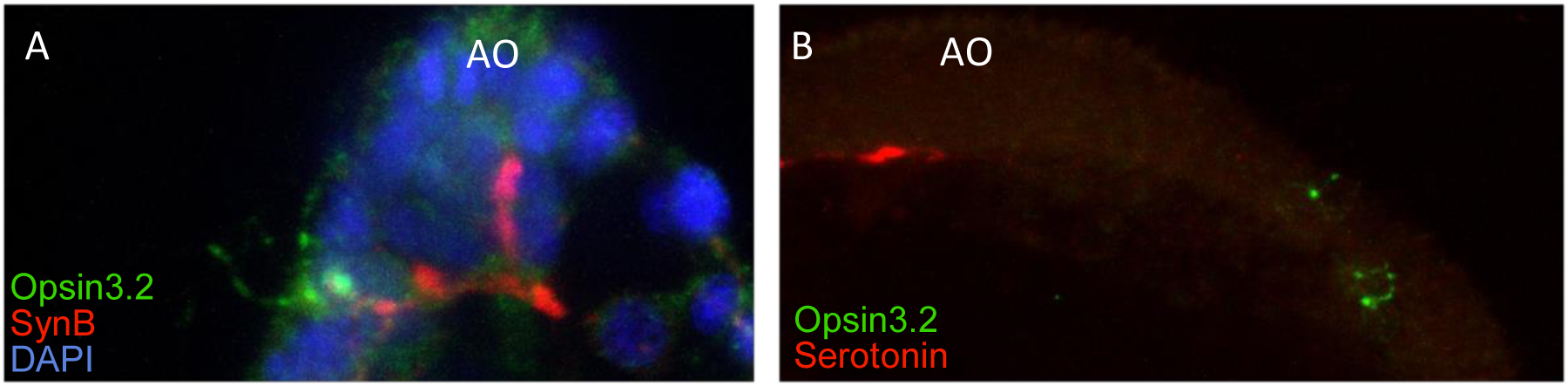
Expression of synaptotagmin but not serotonin in photoreceptors. **(A)** Co-localization of Opsin3.2 and SynaptotagminB. Confocal projection from a left lateral perspective showing the projection of a SynaptotagminB containing axon into the basal neuropil of the Apical Organ (AO). **(B)** Co-immunostaining of Opsin3.2 and Serotonin indicates that PRCs do not contain serotonin. The apical organ is known to include serotonergic neurons on the dorsal margin, and the photoreceptor cells project axons into the apical organ, but do not contain Serotonin.

**Fig. S4.**
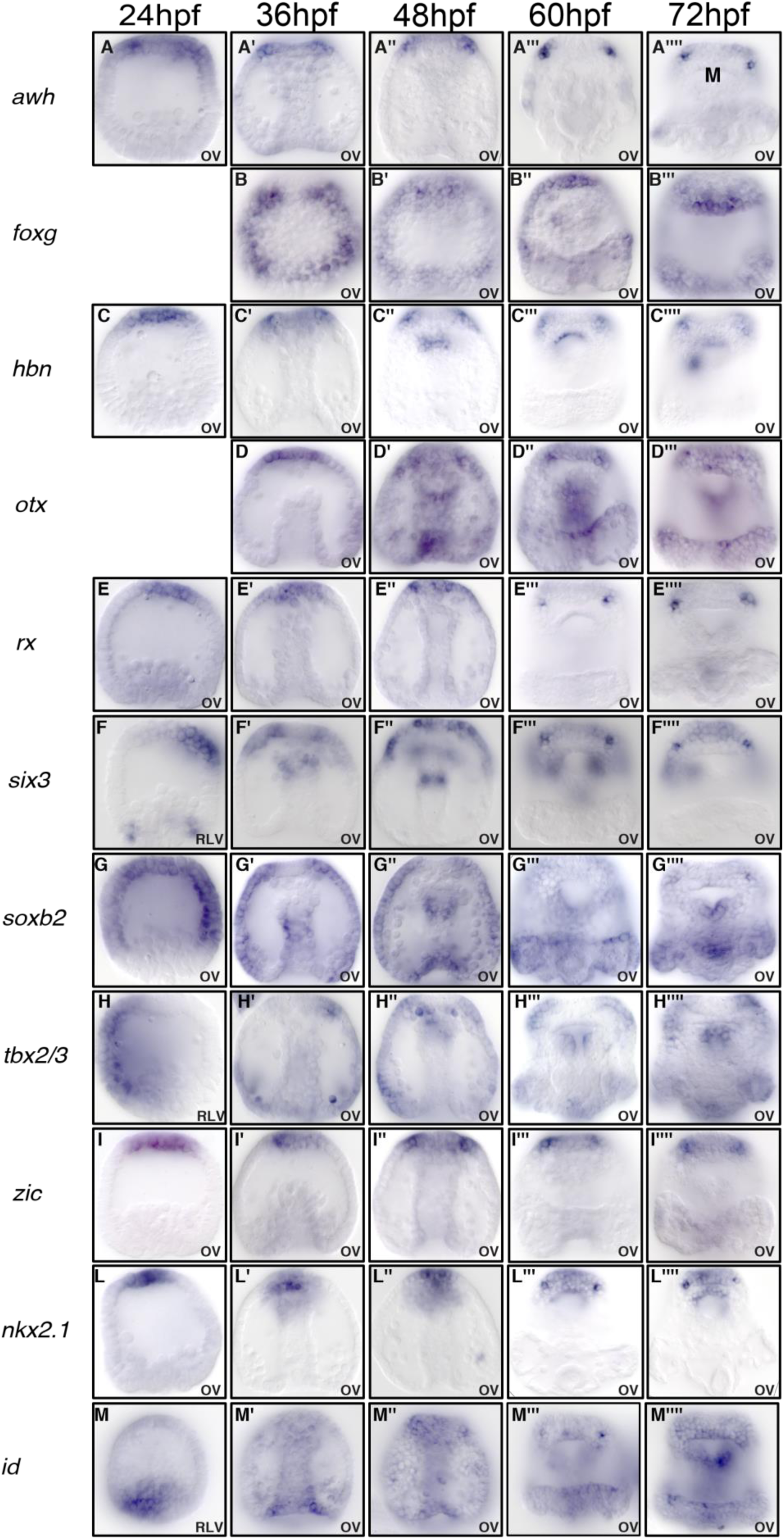
Developmental time course of spatial expression of regulatory genes expressed in photoreceptors. Shown are images of embryos stained by WMISH detecting the expression of *awh* **(A)**, *foxg* **(B)**, *hbn* **(C)**, *otx* **(D)**, *rx* **(E)**, *six3* **(F)**, *soxb2* **(G)**, *tbx2/3* **(H)**, *zic* **(I)**, *nkx2.1* **(L)**, and *id* **(M)**, at 24-72h. OV, oral view; RLV, right lateral view.

**Fig. S5.**
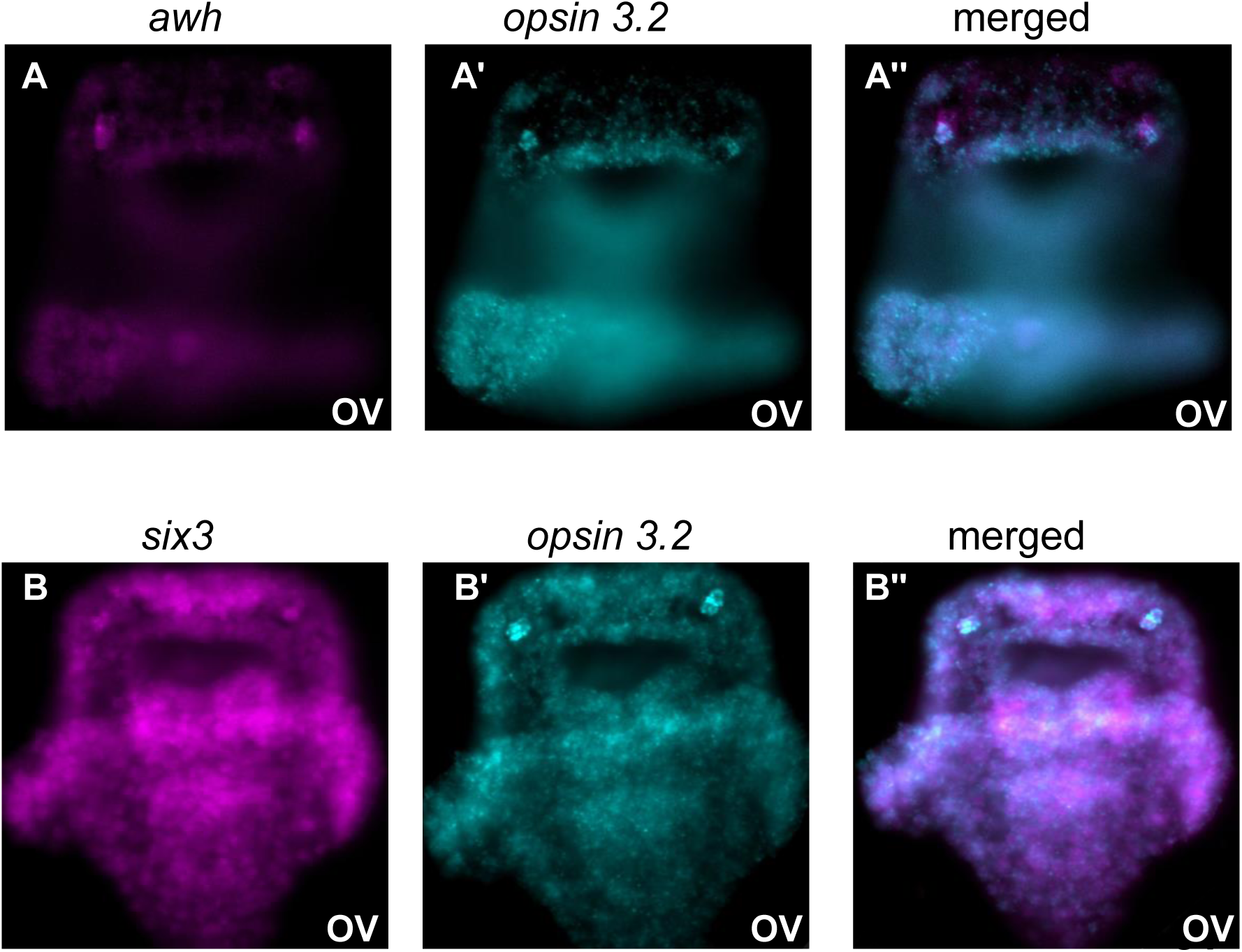
Co-expression of regulatory genes with *opsin3.2* in photoreceptors. Double fluorescent WMISH of opsin3.2 and (A) awh and (B) six3, confirming expression in photoreceptors of 72h sea urchin larvae. OV, oral view.

**Fig. S6.**
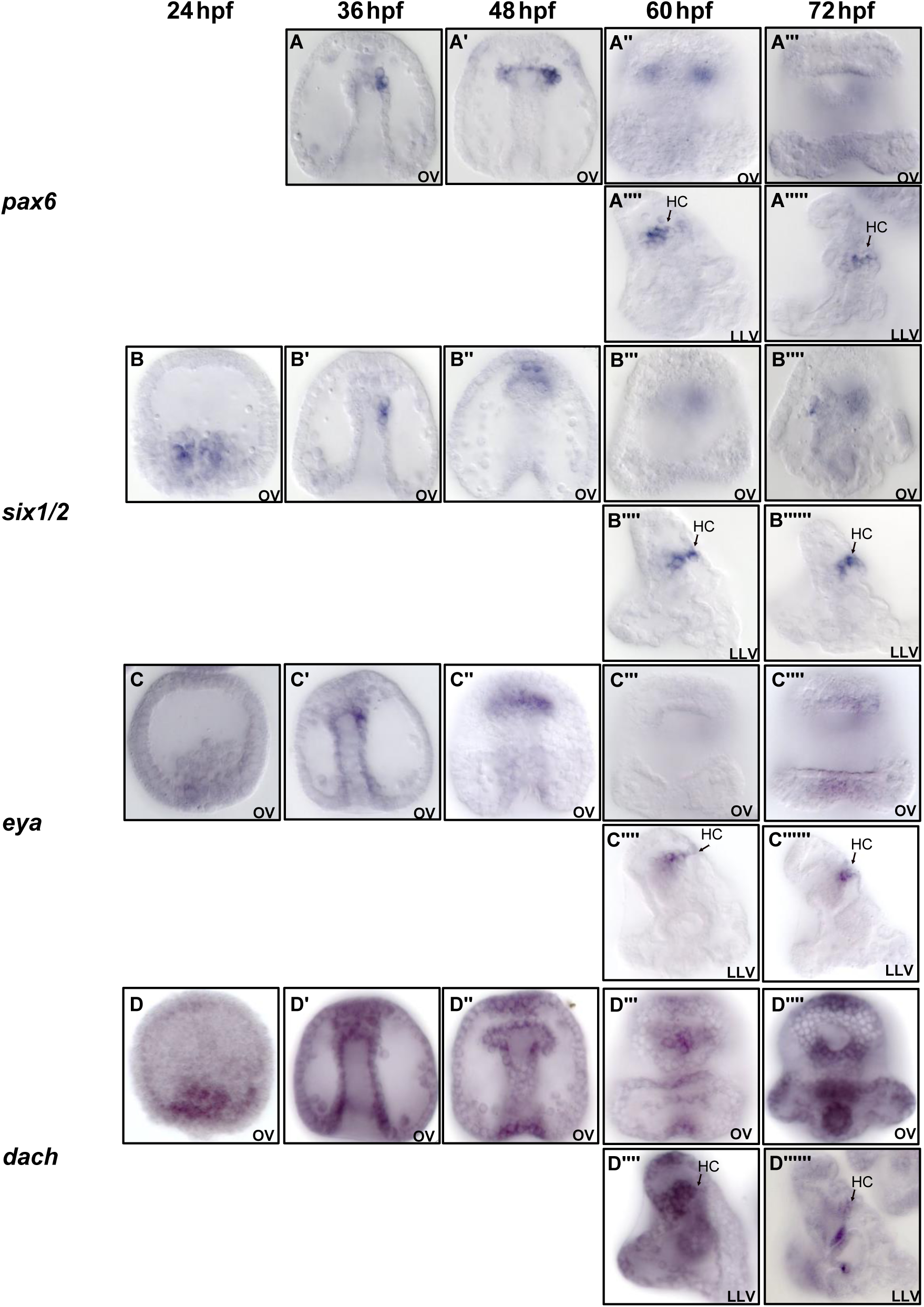
Developmental spatial expression of PSED regulatory genes in the hyrdorpore canal. Images of embryos stained by WMISH for expression of **(A)** *pax6*, **(B)** *six1/2*, **(C)** *eya*, and **(D)** *dach* at 24-72h. OV, oral view; LLV left lateral view; HC, hydropore canal.

**Table S1.**
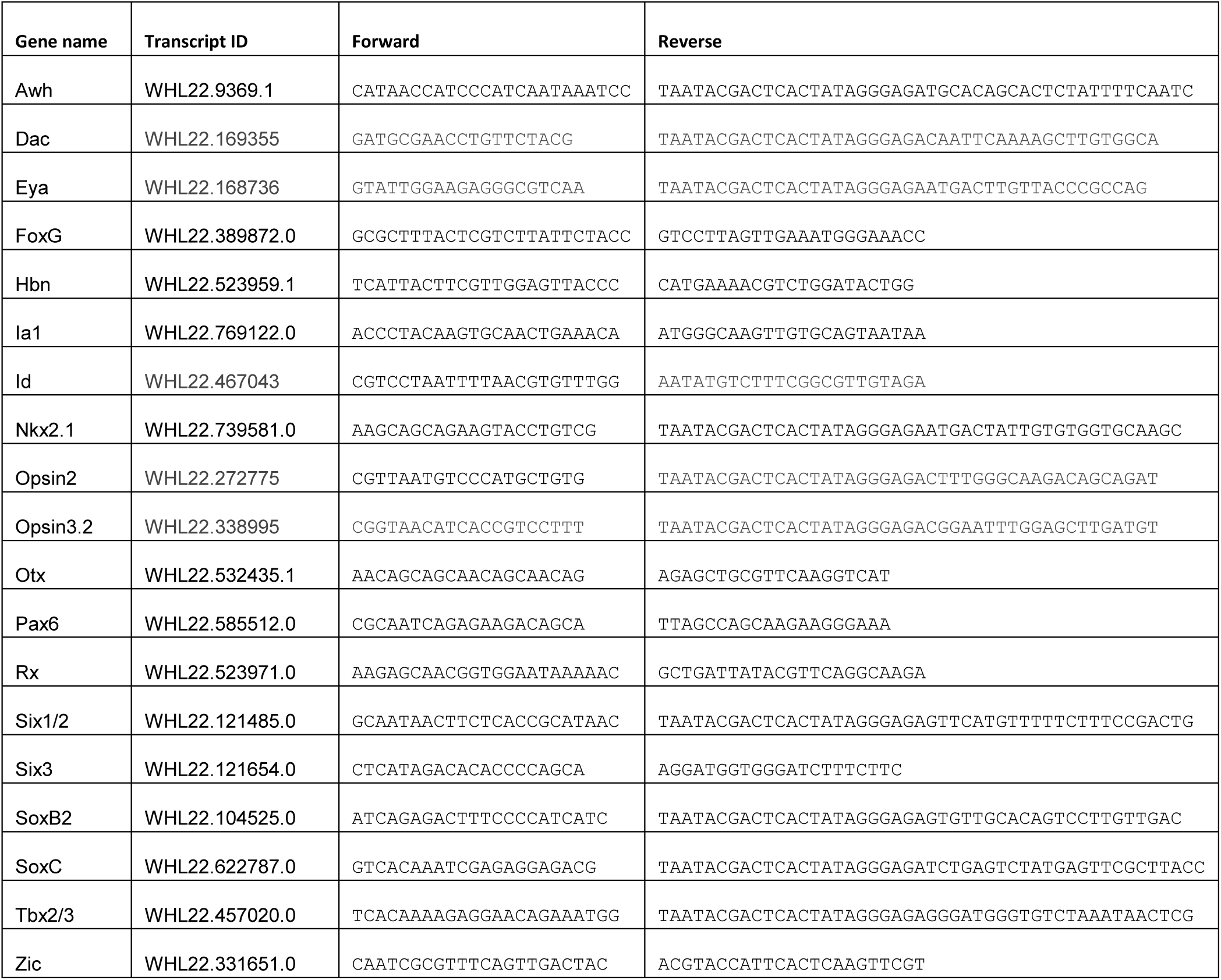
Primers used for generation of WMISH probes. The transcript ID refers to the transcriptome-based gene model defined in (*57*). Gene sequences can be found on Echinobase (http://www.echinobase.org/Echinobase/).

## References

1. Peter, I.S., and Davidson, E.H. (2015). Genomic Control Process, Development and Evolution, (Academic Press/Elsevier).

2. Arendt, D., Musser, J.M., Baker, C.V., Bergman, A., Cepko, C., Erwin, D.H., Pavlicev, M., Schlosser, G., Widder, S., Laubichler, M.D., et al. (2016). The origin and evolution of cell types. Nat Rev Genet 17, 744–757.

3. Peter, I.S., and Davidson, E.H. (2011). Evolution of gene regulatory networks controlling body plan development. Cell 144, 970–985.

4. Arendt, D. (2003). Evolution of eyes and photoreceptor cell types. Int J Dev Biol 47, 563–571.

5. Fain, G.L., Hardie, R., and Laughlin, S.B. (2010). Phototransduction and the evolution of photoreceptors. Current Biology 20, R114–124.

6. Arendt, D., Tessmar-Raible, K., Snyman, H., Dorresteijn, A.W., and Wittbrodt, J. (2004). Ciliary photoreceptors with a vertebrate-type opsin in an invertebrate brain. Science 306, 869–871.

7. Kozmik, Z., Ruzickova, J., Jonasova, K., Matsumoto, Y., Vopalensky, P., Kozmikova, I., Strnad, H., Kawamura, S., Piatigorsky, J., Paces, V., et al. (2008). Assembly of the cnidarian camera-type eye from vertebrate-like components. Proc Natl Acad Sci U S A 105, 8989–8993.

8. Vopalensky, P., Pergner, J., Liegertova, M., Benito-Gutierrez, E., Arendt, D., and Kozmik, Z. (2012). Molecular analysis of the amphioxus frontal eye unravels the evolutionary origin of the retina and pigment cells of the vertebrate eye. Proc Natl Acad Sci U S A 109, 15383–15388.

9. Koyanagi, M., Kubokawa, K., Tsukamoto, H., Shichida, Y., and Terakita, A. (2005). Cephalochordate melanopsin: evolutionary linkage between invertebrate visual cells and vertebrate photosensitive retinal ganglion cells. Curr Biol 15, 1065–1069.

10. Ullrich-Lüter, E.M., Dupont, S., Arboleda, E., Hausen, H., and Arnone, M.I. (2011). Unique system of photoreceptors in sea urchin tube feet. Proc Natl Acad Sci U S A 108, 8367–8372.

11. Arendt, D., and Wittbrodt, J. (2001). Reconstructing the eyes of Urbilateria. Philos Trans R Soc Lond B Biol Sci 356, 1545–1563.

12. Lamb, T.D. (2013). Evolution of phototransduction, vertebrate photoreceptors and retina. Prog Retin Eye Res 36, 52–119.

13. Musser, J.M., and Arendt, D. (2017). Loss and gain of cone types in vertebrate ciliary photoreceptor evolution. Dev Biol 431, 26–35.

14. Bailey, T.J., El-Hodiri, H., Zhang, L., Shah, R., Mathers, P.H., and Jamrich, M. (2004). Regulation of vertebrate eye development by Rx genes. Int J Dev Biol 48, 761–770.

15. Corbo, J.C., Lawrence, K.A., Karlstetter, M., Myers, C.A., Abdelaziz, M., Dirkes, W., Weigelt, K., Seifert, M., Benes, V., Fritsche, L.G., et al. (2010). CRX ChIP-seq reveals the cis-regulatory architecture of mouse photoreceptors. Genome Res.

16. Hennig, A.K., Peng, G.H., and Chen, S. (2008). Regulation of photoreceptor gene expression by Crx-associated transcription factor network. Brain Res. 1192, 114–133.

17. Mathers, P.H., Grinberg, A., Mahon, K.A., and Jamrich, M. (1997). The Rx homeobox gene is essential for vertebrate eye development. Nature 387, 603–607.

18. Nelson, S.M., Park, L., and Stenkamp, D.L. (2009). Retinal homeobox 1 is required for retinal neurogenesis and photoreceptor differentiation in embryonic zebrafish. Dev Biol 328, 24–39.

19. Rodgers, H.M., Huffman, V.J., Voronina, V.A., Lewandoski, M., and Mathers, P.H. (2018). The role of the Rx homeobox gene in retinal progenitor proliferation and cell fate specification. Mech Dev 151, 18–29.

20. Zhang, L., Mathers, P.H., and Jamrich, M. (2000). Function of Rx, but not Pax6, is essential for the formation of retinal progenitor cells in mice. Genesis 28, 135–142.

21. D’Aniello, S., D’Aniello, E., Locascio, A., Memoli, A., Corrado, M., Russo, M.T., Aniello, F., Fucci, L., Brown, E.R., and Branno, M. (2006). The ascidian homolog of the vertebrate homeobox gene Rx is essential for ocellus development and function. Differentiation 74, 222–234.

22. Zuber, M.E., Gestri, G., Viczian, A.S., Barsacchi, G., and Harris, W.A. (2003). Specification of the vertebrate eye by a network of eye field transcription factors. Development (Cambridge, England) 130, 5155–5167.

23. Pan, Y., Martinez-De Luna, R.I., Lou, C.H., Nekkalapudi, S., Kelly, L.E., Sater, A.K., and El-Hodiri, H.M. (2010). Regulation of photoreceptor gene expression by the retinal homeobox (Rx) gene product. Dev Biol 339, 494–506.

24. Tu, Q., Cameron, R.A., and Davidson, E.H. (2014). Quantitative developmental transcriptomes of the sea urchin Strongylocentrotus purpuratus. Dev Biol 385, 160–167.

25. Valero-Gracia, A., Petrone, L., Oliveri, P., Nilsson, D.-E., and Arnone, M.I. (2016). Non-directional photoreceptors in the pluteus of Strongylocentrotus purpuratus. Frontiers in Ecology and Evolution 4, 127.

26. Feuda, R., Hamilton, S.C., McInerney, J.O., and Pisani, D. (2012). Metazoan opsin evolution reveals a simple route to animal vision. Proc Natl Acad Sci U S A 109, 18868–18872.

27. D’Aniello, S., Delroisse, J., Valero-Gracia, A., Lowe, E.K., Byrne, M., Cannon, J.T., Halanych, K.M., Elphick, M.R., Mallefet, J., Kaul-Strehlow, S., et al. (2015). Opsin evolution in the Ambulacraria. Mar Genomics 24 Pt 2, 177–183.

28. Kojima, D., Terakita, A., Ishikawa, T., Tsukahara, Y., Maeda, A., and Shichida, Y. (1997). A novel Go-mediated phototransduction cascade in scallop visual cells. J Biol Chem 272, 22979–22982.

29. Gühmann, M., Jia, H., Randel, N., Verasztó, C., Bezares-Calderón, L.A., Michiels, N.K., Yokoyama, S., and Jékely, G. (2015). Spectral tuning of phototaxis by a go-opsin in the rhabdomeric eyes of Platynereis. Current Biology 25, 2265–2271.

30. Kreft, M., Krizaj, D., Grilc, S., and Zorec, R. (2003). Properties of exocytotic response in vertebrate photoreceptors. Journal of neurophysiology 90, 218–225.

31. Brose, N., Petrenko, A.G., Sudhof, T.C., and Jahn, R. (1992). Synaptotagmin: a calcium sensor on the synaptic vesicle surface. Science (New York, N.Y 256, 1021–1025.

32. Burke, R.D., Osborne, L., Wang, D., Murabe, N., Yaguchi, S., and Nakajima, Y. (2006). Neuron-specific expression of a synaptotagmin gene in the sea urchin Strongylocentrotus purpuratus. J Comp Neurol 496, 244–251.

33. Nakajima, Y., Kaneko, H., Murray, G., and Burke, R.D. (2004). Divergent patterns of neural development in larval echinoids and asteroids. Evol Dev 6, 95–104.

34. Nilsson, D.E. (2009). The evolution of eyes and visually guided behaviour. Philos Trans R Soc Lond B Biol Sci 364, 2833–2847.

35. Gibson, A.W., and Burke, R.D. (1985). The origin of pigment cells in embryos of the sea urchin Strongylocentrotus purpuratus. Dev Biol 107, 414–419.

36. Burke, R.D., Angerer, L.M., Elphick, M.R., Humphrey, G.W., Yaguchi, S., Kiyama, T., Liang, S., Mu, X., Agca, C., Klein, W.H., et al. (2006). A genomic view of the sea urchin nervous system. Dev Biol 300, 434–460.

37. Luo, Y.J., and Su, Y.H. (2012). Opposing nodal and BMP signals regulate left-right asymmetry in the sea urchin larva. PLoS Biol 10, e1001402.

38. Martik, M.L., and McClay, D.R. (2015). Deployment of a retinal determination gene network drives directed cell migration in the sea urchin embryo. 1–19.

39. Ullrich-Luter, E.M., D’Aniello, S., and Arnone, M.I. (2013). C-opsin expressing photoreceptors in echinoderms. Integr Comp Biol 53, 27–38.

40. Swaroop, A., Kim, D., and Forrest, D. (2010). Transcriptional regulation of photoreceptor development and homeostasis in the mammalian retina. Nat. Rev. Neurosci. 11, 563–576.

41. Passamaneck, Y.J., Furchheim, N., Hejnol, A., Martindale, M.Q., and Lüter, C. (2011). Ciliary photoreceptors in the cerebral eyes of a protostome larva. EvoDevo 2, 6–6.

42. Sotolongo-Lopez, M., Alvarez-Delfin, K., Saade, C.J., Vera, D.L., and Fadool, J.M. (2016). Genetic Dissection of Dual Roles for the Transcription Factor six7 in Photoreceptor Development and Patterning in Zebrafish. PLoS Genet 12, e1005968.

43. Ogawa, Y., Shiraki, T., Kojima, D., and Fukada, Y. (2015). Homeobox transcription factor Six7 governs expression of green opsin genes in zebrafish. Proc Biol Sci 282, 20150659.

44. Manavathi, B., Peng, S., Rayala, S.K., Talukder, A.H., Wang, M.H., Wang, R.A., Balasenthil, S., Agarwal, N., Frishman, L.J., and Kumar, R. (2007). Repression of Six3 by a corepressor regulates rhodopsin expression. Proc Natl Acad Sci U S A 104, 13128–13133.

45. Alvarez-Delfin, K., Morris, A.C., Snelson, C.D., Gamse, J.T., Gupta, T., Marlow, F.L., Mullins, M.C., Burgess, H.A., Granato, M., and Fadool, J.M. (2009). Tbx2b is required for ultraviolet photoreceptor cell specification during zebrafish retinal development. Proc Natl Acad Sci U S A 106, 2023–2028.

46. Luo, J., Uribe, R.A., Hayton, S., Calinescu, A.A., Gross, J.M., and Hitchcock, P.F. (2012). Midkine-A functions upstream of Id2a to regulate cell cycle kinetics in the developing vertebrate retina. Neural Dev 7, 33.

47. Mizeracka, K., DeMaso, C.R., and Cepko, C.L. (2013). Notch1 is required in newly postmitotic cells to inhibit the rod photoreceptor fate. Development (Cambridge, England) 140, 3188–3197.

48. Watabe, Y., Baba, Y., Nakauchi, H., Mizota, A., and Watanabe, S. (2011). The role of Zic family zinc finger transcription factors in the proliferation and differentiation of retinal progenitor cells. Biochem Biophys Res Commun 415, 42–47.

49. Oonuma, K., Tanaka, M., Nishitsuji, K., Kato, Y., Shimai, K., and Kusakabe, T.G. (2016). Revised lineage of larval photoreceptor cells in Ciona reveals archetypal collaboration between neural tube and neural crest in sensory organ formation. Dev Biol 420, 178–185.

50. Kumar, J.P. (2009). The molecular circuitry governing retinal determination. Biochim. Biophys. Acta 1789, 306–314.

51. Quiring, R., Walldorf, U., Kloter, U., and Gehring, W.J. (1994). Homology of the eyeless gene of Drosophila to the Small eye gene in mice and Aniridia in humans. Science (New York, N.Y 265, 785–789.

52. Marquardt, T., Ashery-Padan, R., Andrejewski, N., Scardigli, R., Guillemot, F., and Gruss, P. (2001). Pax6 is required for the multipotent state of retinal progenitor cells. Cell 105, 43–55.

53. Chow, R.L., Altmann, C.R., Lang, R.A., and Hemmati-Brivanlou, A. (1999). Pax6 induces ectopic eyes in a vertebrate. Development (Cambridge, England) 126, 4213–4222.

54. Halder, G., Callaerts, P., and Gehring, W.J. (1995). Induction of ectopic eyes by targeted expression of the eyeless gene in Drosophila. Science (New York, N.Y 267, 1788–1792.

55. Gehring, W.J., and Ikeo, K. (1999). Pax 6: mastering eye morphogenesis and eye evolution. Trends Genet 15, 371–377.

56. Nilsson, D.E. (2004). Eye evolution: a question of genetic promiscuity. Curr Opin Neurobiol 14, 407–414.

57. Irvine, S.Q., Fonseca, V.C., Zompa, M.A., and Antony, R. (2008). Cis-regulatory organization of the Pax6 gene in the ascidian Ciona intestinalis. Dev Biol 317, 649–659.

58. Pichaud, F., Treisman, J., and Desplan, C. (2001). Reinventing a common strategy for patterning the eye. Cell 105, 9–12.

59. Remez, L.A., Onishi, A., Menuchin-Lasowski, Y., Biran, A., Blackshaw, S., Wahlin, K.J., Zack, D.J., and Ashery-Padan, R. (2017). Pax6 is essential for the generation of late-born retinal neurons and for inhibition of photoreceptor-fate during late stages of retinogenesis. Dev Biol 432, 140–150.

60. Vopalensky, P., and Kozmik, Z. (2009). Eye evolution: common use and independent recruitment of genetic components. Philos Trans R Soc Lond B Biol Sci 364, 2819–2832.

61. Kozmik, Z., Holland, N.D., Kreslova, J., Oliveri, D., Schubert, M., Jonasova, K., Holland, L.Z., Pestarino, M., Benes, V., and Candiani, S. (2007). Pax-Six-Eya-Dach network during amphioxus development: conservation in vitro but context specificity in vivo. Dev Biol 306, 143–159.

62. Heanue, T.A., Reshef, R., Davis, R.J., Mardon, G., Oliver, G., Tomarev, S., Lassar, A.B., and Tabin, C.J. (1999). Synergistic regulation of vertebrate muscle development by Dach2, Eya2, and Six1, homologs of genes required for Drosophila eye formation. Genes Dev 13, 3231–3243.

63. Nilsson, D.E. (1996). Eye ancestry: old genes for new eyes. Curr Biol 6, 39–42.

64. Putnam, N.H., Butts, T., Ferrier, D.E., Furlong, R.F., Hellsten, U., Kawashima, T., Robinson-Rechavi, M., Shoguchi, E., Terry, A., Yu, J.K., et al. (2008). The amphioxus genome and the evolution of the chordate karyotype. Nature 453, 1064–1071.

65. Huang, S., Chen, Z., Yan, X., Yu, T., Huang, G., Yan, Q., Pontarotti, P.A., Zhao, H., Li, J., Yang, P., et al. (2014). Decelerated genome evolution in modern vertebrates revealed by analysis of multiple lancelet genomes. Nat Commun 5, 5896.

66. Dehal, P., Satou, Y., Campbell, R.K., Chapman, J., Degnan, B., De Tomaso, A., Davidson, B., Di Gregorio, A., Gelpke, M., Goodstein, D.M., et al. (2002). The draft genome of Ciona intestinalis: insights into chordate and vertebrate origins. Science (New York, N.Y 298, 2157–2167.

67. Small, K.S., Brudno, M., Hill, M.M., and Sidow, A. (2007). A haplome alignment and reference sequence of the highly polymorphic Ciona savignyi genome. Genome Biol 8, R41.

68. Altschul, S.F., Gish, W., Miller, W., Myers, E.W., and Lipman, D.J. (1990). Basic local alignment search tool. J Mol Biol 215, 403–410.

69. Quevillon, E., Silventoinen, V., Pillai, S., Harte, N., Mulder, N., Apweiler, R., and Lopez, R. (2005). InterProScan: protein domains identifier. Nucleic Acids Res 33, W116–120.

70. Katoh, K., Misawa, K., Kuma, K., and Miyata, T. (2002). MAFFT: a novel method for rapid multiple sequence alignment based on fast Fourier transform. Nucleic Acids Res 30, 3059–3066.

71. Le, S.Q., and Gascuel, O. (2008). An improved general amino acid replacement matrix. Mol Biol Evol 25, 1307–1320.

72. Nguyen, L.T., Schmidt, H.A., von Haeseler, A., and Minh, B.Q. (2015). IQ-TREE: a fast and effective stochastic algorithm for estimating maximum-likelihood phylogenies. Mol Biol Evol 32, 268–274.

73. Minh, B.Q., Nguyen, M.A., and von Haeseler, A. (2013). Ultrafast approximation for phylogenetic bootstrap. Mol Biol Evol 30, 1188–1195.

74. Anisimova, M., Gil, M., Dufayard, J.F., Dessimoz, C., and Gascuel, O. (2011). Survey of branch support methods demonstrates accuracy, power, and robustness of fast likelihood-based approximation schemes. Syst Biol 60, 685–699.

75. Lartillot, N., Lepage, T., and Blanquart, S. (2009). PhyloBayes 3: a Bayesian software package for phylogenetic reconstruction and molecular dating. Bioinformatics 25, 2286–2288.

76. Huerta-Cepas, J., Szklarczyk, D., Forslund, K., Cook, H., Heller, D., Walter, M.C., Rattei, T., Mende, D.R., Sunagawa, S., Kuhn, M., et al. (2016). eggNOG 4.5: a hierarchical orthology framework with improved functional annotations for eukaryotic, prokaryotic and viral sequences. Nucleic Acids Res 44, D286–293.

77. Ransick, A., Ernst, S., Britten, R.J., and Davidson, E.H. (1993). Whole mount in situ hybridization shows Endo 16 to be a marker for the vegetal plate territory in sea urchin embryos. Mechanisms of development 42, 117–124.

78. Chen, J.H., Luo, Y.J., and Su, Y.H. (2011). The dynamic gene expression patterns of transcription factors constituting the sea urchin aboral ectoderm gene regulatory network. Dev Dyn 240, 250–260.

79. Garner, S., Zysk, I., Byrne, G., Kramer, M., Moller, D., Taylor, V., and Burke, R.D. (2016). Neurogenesis in sea urchin embryos and the diversity of deuterostome neurogenic mechanisms. Development 143, 286–297.

80. Vielkind, U., and Swierenga, S.H. (1989). A simple fixation procedure for immunofluorescent detection of different cytoskeletal components within the same cell. Histochemistry 91, 81–88.

## Supplemental References

1. Tu, Q., Cameron, R.A., and Davidson, E.H. (2014). Quantitative developmental transcriptomes of the sea urchin Strongylocentrotus purpuratus. Dev Biol 385, 160–167.

2. Feuda, R., Hamilton, S.C., McInerney, J.O., and Pisani, D. (2012). Metazoan opsin evolution reveals a simple route to animal vision. Proc Natl Acad Sci U S A 109, 18868–18872.

3. Guhmann, M., Jia, H., Randel, N., Veraszto, C., Bezares-Calderon, L.A., Michiels, N.K., Yokoyama, S., and Jekely, G. (2015). Spectral Tuning of Phototaxis by a Go-Opsin in the Rhabdomeric Eyes of Platynereis. Curr Biol 25, 2265–2271.

